# Colour preference and constancy in the giant Asian honey bee *Apis dorsata*

**DOI:** 10.64898/2026.01.21.700526

**Authors:** R Sudeep, Sachin Bhaskar, Hema Somanathan

**Affiliations:** School of Biology, Indian Institute of Science Education and Research Thiruvananthapuram, Kerala, India

## Abstract

Tropical pollinators forage in environments where floral resources vary in space and time, requiring flexible strategies to optimise foraging efficiency. One such strategy, floral constancy - the temporary restriction to a single flower type - strongly influences foraging success and plant-pollinator interactions. We aimed to: (1) quantify spontaneous colour preferences and constancy in the Asian giant honeybee *Apis dorsata*, (2) test whether reward concentration modulates these preferences, (3) evaluate how quickly learned associations override spontaneous biases, (4) determine whether bees can use multiple colour associations simultaneously, and (5) assess whether local floral spectral patterns correlate with bee preferences.

Bees trained to a neutral UV-grey stimulus showed a strong spontaneous preference and high constancy to blue, revealing a robust short-wavelength bias. Crucially, the strength of this spontaneous bias depended on reward concentration; Low-reward conditions elicited strong blue constancy, whereas high-reward conditions weakened it, demonstrating that reward expectation shapes spontaneous colour choices. This bias was flexible. When bees learned that yellow was rewarding they switched their preferences. Bees sequentially trained to both colours visited blue and yellow, showing no overall bias, or effect of the last-trained colour, indicating that recent experiences disrupt colour-specific constancy and generate largely random foraging choices. Bees were capable of learning and retaining two colours simultaneously, effectively suppressing the influence of spontaneous preferences. Finally, analysis of the community’s floral spectral distribution revealed a strong dominance of short-wavelength flowers, suggesting that long-term selection by the local floral environment may underlie the spontaneous blue preference observed in *A. dorsata*.

## Introduction

Foraging pollinators encounter a wide variety of flowers while searching for resources, yet many display temporary fidelity to a single flower species during a foraging bout while avoiding other co-occurring flowers (Amaya-Márquez, 2009; Grant, 1950; Grüter & Ratnieks, 2011; Waser, 1986). This behaviour, known as floral constancy, has been documented in diverse pollinator groups, including bees, birds, butterflies, and flies (Goulson & Cory, 1993; Goulson & Wright, 1998; Grant, 1950; Lewis, 1986; Schmid et al., 2016; Shrotri et al., 2024; Slaa et al., 1998; Somanathan et al., 2019; Thomson, 1981; Waser, 1986; Weiss, 2001; Wells & Rathore, 1994). Floral constancy has important consequences for plant fitness by increasing the probability of conspecific pollen transfer. The adaptive value of constancy in pollinators has been discussed in relation to neuro-sensory constraints in recalling diverse floral traits (Gegear & Laverty, 2005; Goulson, 2000; Goulson et al., 1997; Hill et al., 2001; Ishii, 2005; Ishii & Masuda, 2014). Such constraints are thought to be alleviated by specialising on one floral type, thereby enhancing pollinator efficiency in searching for, handling, and utilising rewards (Amaya-Márquez, 2009; Chittka et al., 1999; Grüter et al., 2011; Grüter & Ratnieks, 2011; Hill et al., 2001; Takagi & Ohashi, 2025).

Pollinators show varying degrees of floral constancy depending on factors such as colour, odour, reward quality and quantity, and the distribution and abundance of flowers (Baude et al., 2011; Bruninga-Socolar et al., 2022; Gegear & Laverty, 2004, 2005; Hill et al., 1997; Ishii & Kadoya, 2016; Nityananda & Chittka, 2021; Slaa et al., 2003). Colour is especially important for visually guided pollinators like bees, many of which show innate (or spontaneous) biases for particular spectral wavelengths (Chittka & Menzel, 1992; van der Kooi et al., 2019). For example, *Apis mellifera, Bombus terrestris*, and *Tetragonula iridipennis* prefer short-wavelength colours such as UV–blue, blue, and violet (Balamurali et al., 2018; Giurfa et al., 1995; Gumbert, 2000), whereas *Apis cerana* is biased toward longer wavelengths such as blue-green and lime-yellow (Balamurali et al., 2018). These biases likely reflect evolutionary associations with reliably rewarding flowers. However, colour preferences are highly plastic and pollinators can rapidly learn and modify their choices based on the local floral environment and prior experience (Chittka & Raine, 2006; Dyer & Chittka, 2004; Giurfa et al., 1995; Kinoshita et al., 2017; Menzel & Shmida, 1993; Raine & Chittka, 2007b). Experience with alternative colours can override innate tendencies (Giurfa et al., 1995; Heinrich et al., 1977; Menzel, 1985), facilitating the exploration of a broader range of flowers which is particularly important in dynamic floral environments. How innate and learned preferences interact remains debated. One view argues that early colour learning rapidly replaces innate biases, while another holds that spontaneous preferences remain stable throughout a forager’s lifetime and continue to influence decisions in unfamiliar contexts. Both perspectives suggest a close relationship between innate preferences, the formation and recall of colour memories, although the underlying mechanisms are not yet understood.

These dynamics of innate bias and learning form the basis for understanding how pollinators maintain constancy when foraging among diverse floral resources. Consequently, floral constancy has been studied across multiple systems by using various approaches to evaluate how these behaviours manifest under natural and experimental conditions. Floral constancy has been examined using field observations by following the movement of foraging pollinators (Bruninga-Socolar et al., 2022; De Jager et al., 2011; Raine & Chittka, 2007a; van der Niet et al., 2020; Waser, 1986; Wilson & Stine, 1996), from pollen loads on pollinators bodies (Raine & Chittka, 2005b; Shrotri et al., 2024; Somanathan et al., 2020; Yourstone et al., 2023), from laboratory studies of pollinator choices using artificial flower arrays (Grüter et al., 2011; Heinrich et al., 1977; Ishii & Masuda, 2014; Takagi & Ohashi, 2025; Wells et al., 1992; Wells & Rathore, 1994), and more recently using agent-based modelling approaches (Hayes & Grüter, 2023). Unsurprisingly, tropical honeybee species are little studied apart from a few on the Eastern honeybee *Apis cerana* (Wells et al., 1992; Wells & Rathore, 1994). Given species-specific variation even among closely related bees, understanding pollinator behaviour in tropical systems is essential (Grüter & Ratnieks, 2011; Somanathan, 2024; Somanathan & Balamurali, 2023). Species level differences may reflect the distinct ecological contexts in which species have evolved. Factors such as floral phenology and competition can influence pollinator preferences and constancy (Ogilvie & Forrest, 2017). Tropical habitats are also marked by high richness of plants and pollinators (Ollerton et al., 2006; Somanathan, 2024; Vizentin-Bugoni et al., 2018), greater phenological diversity, and a broader range of reproductive strategies than known from temperate regions (Morellato et al., 2013; Newstrom et al., 1994). Therefore it is critical to examine foraging behaviours in tropical honey bees in their native habitats. Examining floral constancy in tropical honeybees provides an opportunity to uncover how pollinator foraging strategies are shaped under the complex ecological pressures of tropical plant–pollinator communities.

Despite this extensive body of research, the interplay between innate (or spontaneous) colour preferences, learning, and the motivational state of bees remain poorly understood. How these interacting factors shape floral constancy in foraging situations has received remarkably little direct attention. In this study, we investigate floral constancy to colour in naturally occurring wild colonies of the giant Asian honeybee *Apis dorsata*, a common tropical open-nesting species found throughout South and Southeast Asia. Unlike the cavity-nesting *A. mellifera* and *A. cerana*, *A. dorsata* cannot be domesticated or maintained in managed hives, and therefore all experiments must be conducted on bees recruited from natural colonies. We examine the spontaneous colour preferences of wild foragers (whose prior experience remains unknown) after first removing any prior colour-specific experience by training them to a neutral UV-grey stimulus. Previous work has shown that such neutral-stimulus training effectively erases learned colour associations, allowing bees to revert to their innate colour preferences (Balamurali et al., 2018; Gumbert, 2000). Using blue and yellow dimorphic artificial flower arrays provisioned with sugar solution as reward, we addressed the following questions: (1) Does *A. dorsata* exhibit spontaneous floral constancy to colour? (2) Is spontaneous floral constancy modulated by the quality (concentration) of the sugar reward? (3) Is spontaneous constancy modulated by recent foraging experience with colour? Finally, a prevalent hypothesis suggests that that spontaneous colour preference for blue in bees (Giurfa et al., 1995; Gumbert, 2000) have evolved in response to the dominant floral colour in plant communities. Therefore, we asked if: (4) Floral spectral properties of the surrounding floral community can explain the spontaneous colour preference in *A. dorsata*?

## Methods

### Study species and site

We conducted our experiments using foragers from wild *Apis dorsata* colonies nesting on the roof overhangs of the Biological Science Building at the Indian Institute of Science Education and Research Thiruvananthapuram (IISER TVM; 08°40’55.5398“ N, 077°08’08.4215” W) (Figure.1A; Supplementary figure 1). During the study period from December 2020 to August 2022, colonies underwent episodes of swarming and migration, so the number of active colonies fluctuated from a maximum of 16 to a minimum of three.

**Figure 1:**
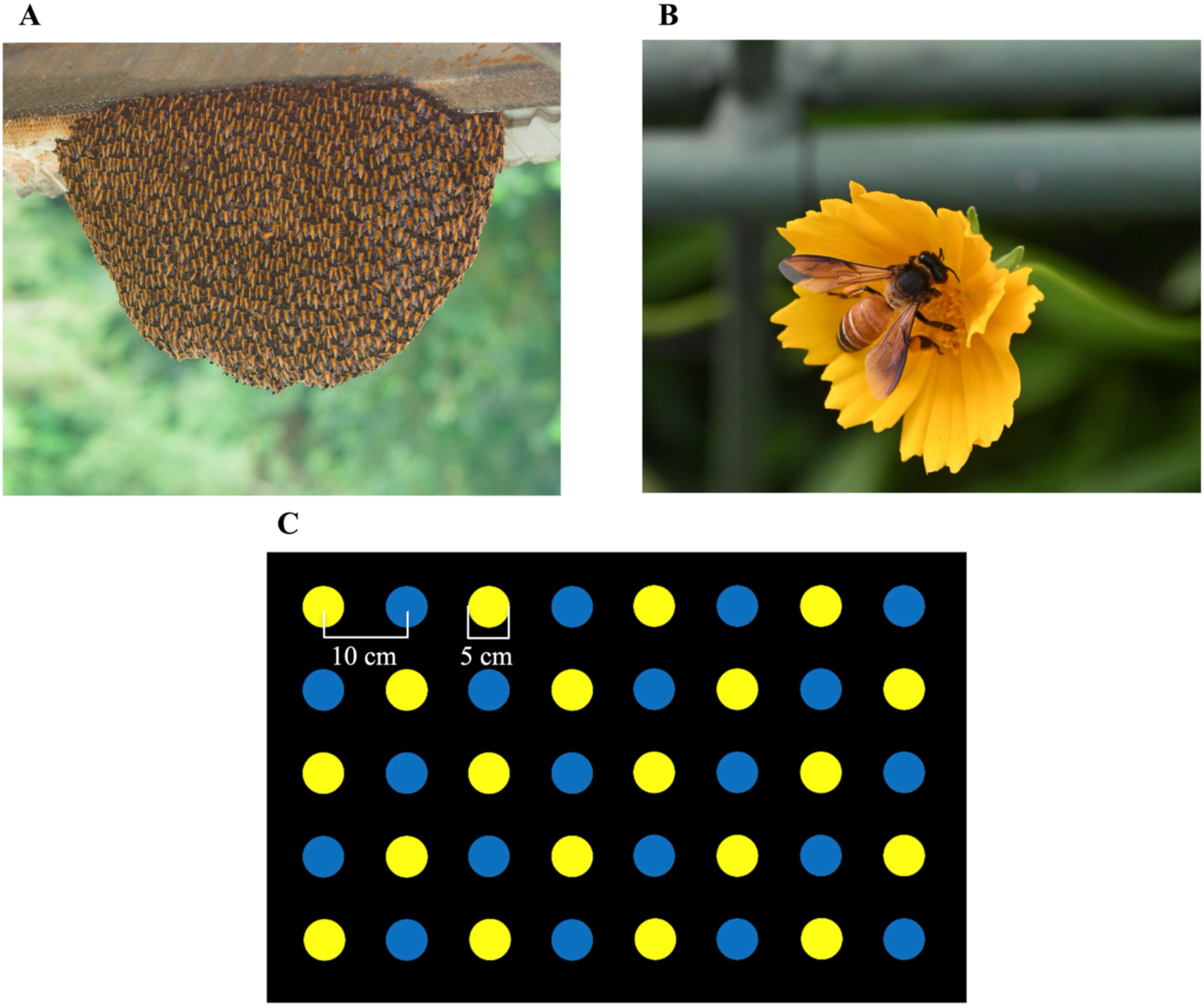
**A)** *Apis dorsata* colony on the roof overhangs of the Biological Sciences Building at IISER Thiruvananthapuram **B)** *A. dorsata* on a flower (*Cosmos sulphureus*) **C)** A schematic of the blue and yellow test array.

Foragers were first trained to a black feeder containing sugar solution. After enough bees had been recruited, the feeder was gradually moved to an experimental arena located on the roof terrace about 5-10 m from the hives. The arena consisted of an aluminium cage (1m x 1m x 0.75m) with nylon mesh sides and a plexiglass sheet roof. A video camera (GoPro Hero 7 Black) was mounted at the top of the cage to record the experiments. Bees entered the arena through a clear plexiglass tunnel equipped with shutters that allowed us to control the number of bees entering the setup (Supplementary figure 2).

### Stimuli

Matte finish vinyl stickers (LG Hausys) pasted on clear 5cm diameter acrylic disks were used as training and test stimuli in the experiments. UV-grey stimuli made from matte finish UV reflecting cardboard sheets were pasted on the clear acrylic disks. Matte black vinyl stickers were used as the background on which the stimuli were presented in all experiments. The spectral reflectance of the stimuli was measured using a spectrophotometer (Ocean Optics Ocean-HDX-UV-vis spectrometer, supplemented by a PX-2 pulsed Xenon light source; supplementary figure 3A). The colour loci of the stimuli were modelled in the hexagonal colour space (Chittka, 1992) using the *Pavo* package (Maia et al., 2013) in R, version 4.5.1 (R Core Team, 2025), to ensure that the colours used in the experiments were discriminable by bees from the background achromatically and chromatically (Supplementary table T1).

All experiments consisted of three phases: a pre-training phase, a training phase, and a test phase. The pre-training phase was identical in all experiments during which bees that were recruited into the arena were presented with a pair of clear acrylic disks at the centre of which sugar solution was provided *ad libitum*. While they were feeding, bees were individually marked using acrylic paint (Camel Fabrica, Camlin Kokuyo, India) to identify individuals. Following this, marked individuals were allowed to forage from a different set of four clear acrylic disks for three consecutive foraging bouts. A bout consisted of all the stimuli choices of a bee from the time of entering to exiting the arena. The positions of the disks were changed frequently to encourage bees to explore all the disks and to avoid any positional bias.

#### I. Spontaneous constancy to colour under different training and test reward concentrations

To examine spontaneous constancy and whether it is modulated by reward quality (sugar concentration), we performed three experiments (a-c below) in which the reward concentration varied during training and testing phases.

In the following three experiments (a-c), we followed a common protocol where bees were trained to the arena and fed on sugar solution provided in four UV-grey stimuli. The UV-grey stimulus serves as a neutral achromatic stimulus that is shown to erase learned associations and prompts the bees to rely on spontaneously preferred colour while making foraging decisions (Balamurali et al., 2018; Gumbert, 2000). The stimuli positions were changed during training to encourage exploration and avoid positional learning. Each UV-grey stimulus contained 30µl of sugar solution and was replenished after a bee moved to feed from another stimulus. A training bout was considered successful if the marked bee fed from multiple stimuli before leaving the arena. Since the bees fed from 3 to 4 stimuli during each training bout and each training stimulus contained 30µl of sugar solution, the average crop capacity of bees was estimated to be between 90µl and 120µl. After the completion of four training bouts, bees were tested on their next return to the arena to an alternative array of 20 blue and 20 yellow stimuli with an inter-stimulus distance (centre to centre) of 10 cm (Figure 1C) that contained 5µl of sugar solution. Providing a small volume of sugar encouraged bees to visit multiple stimuli during a test before they reached satiation and exited the arena. Only bees that completed training and returned within two minutes were included in the test, in order to ensure that they did not visit other foraging resources in the habitat. A single bee was allowed to enter the arena at a given time during training and testing. Landings on stimuli were considered as choices in tests. Tested bees were euthanised to avoid pseudoreplication. Tests were recorded (1920 x 1080 pixel resolution at 30 fps) using a video camera (GoPro Hero 7 Black), and were later manually analysed frame-by-frame using QuickTime Player (version 10.5). The three experimental reward conditions (a-c) that the bees experienced are described below.

##### a) Training and testing with low reward concentration (10%)

This experiment was designed to test for constancy to colour under low-reward conditions, after training to a neutral colour stimuli. Training to neutral colour is known to induce a return to spontaneous colour preferences in previous studies (Balamurali et al., 2018; Gumbert, 2000). Bees were first trained to feed from four UV-grey stimuli, each provisioned with 10% sugar solution (w/w). After completing training, each bee was tested in the blue-yellow array where all stimuli contained 10% sugar solution (n = 22, Fig 1C).

##### b) Training with high reward concentration (30%) and testing with low reward concentration (10%)

This experiment was designed to test whether training to a higher concentration reward alters the colour preference in tests when the bees are presented with low-concentration reward. During training, the UV-grey stimuli were provisioned with 30% sugar solution (n = 21). Each bee completed four training bouts with the UV-grey stimulus before it was tested (as mentioned above) using the blue-yellow stimuli containing 10% sugar solution.

##### c) Training and testing with high reward concentration (30%)

In this experiment we examine colour choices of bees when both training and testing were carried out under consistently high-reward conditions during training and testing. Bees were first trained on a neutral UV-grey stimulus as in the previous two experiments, each provisioned with 30% sugar solution. Each bee was tested using the blue and yellow array, each of which contained 30% sugar solution.

#### II. Effect of recent foraging experience on constancy

To examine if spontaneous constancy to blue colour that was observed in the above experiments was affected by recent foraging experience, we performed two experiments (*d* and *e*) in which bees were trained to either the less preferred yellow or alternatively on blue and yellow.

##### d) Training to yellow

A set of bees were trained on the less preferred yellow (n=22) containing 30µl of 10% sugar solution and then tested with the blue-yellow stimulus array each of which was rewarded with 5µl of 10% sugar solution.

##### e) Alternate training to blue and yellow

Next, we addressed the effect of recent foraging experience on colour preference and constancy. Bees were trained in an alternating sequence to both blue and yellow stimuli provided with 30µl of 10% sugar solution. All bees were trained on each colour alternatively; in one set of bees the first training was to blue and the last training was to yellow (*blue-yellow-blue-yellow-blue-yellow-blue-yellow,* n = 20) and another set of bees was trained in the opposite sequence (*yellow-blue-yellow-blue-yellow-blue-yellow-blue,* n = 20). These bees were tested on an array where both the colours had 5µl of 10% sugar solution.

### Floral community spectra

We measured the floral reflectance spectra from 123 plant species within a 2 km radius of the experimental hives, a distance well within the known foraging ranges of *A. dorsata* (Young et al., 2021). Reflectance measurements of petals were taken using a spectrophotometer (Ocean Optics Ocean-HDX-UV-vis spectrometer, supplemented by a PX-2 pulsed Xenon light source), with 3-6 biological replicates per species except for rare species. For each species, one representative spectral curve was used for analysis. To quantify floral contrast against natural backgrounds, which is a key determinant of how bees perceive and detect flowers, we also measured the background leaf spectra for 83 species and used the median leaf spectrum as a standard green background.

Floral spectra were then modelled in the bee colour hexagon (Chittka, 1992), which incorporates both receptor excitation and chromatic contrast relative to a natural background. This allowed us to examine the distribution of bee-perceived colours within the local plant community and assess whether certain spectral regions are overrepresented in the floral community. Plants were classified as native (n = 65) or exotic (n = 58) using the Kew “Plants of the World Online” database (*Plants of the World Online | Kew Science*, 2025) and analysed both separately and jointly. Each species was assigned to one of the six bee-subjective colour categories corresponding to the sectors of the hexagon model (UV, UV-blue, blue, blue-green, green, and UV-green), and then further grouped into short-wavelength (UV, UV-blue, blue) and long-wavelength (blue-green, green, UV-green) categories to evaluate broad spectral patterns in the floral community.

### Statistical analyses

#### Behavioural experiments

All statistical analyses were performed in R (version 4.5.1). In all experiments (*a-e*), the first and total choices made by bees to each colour were obtained by analysing videos frame-by-frame using QuickTime Player (version 10.5). A test of proportions was used to compare the first choices of bees. Wilcoxon tests were performed to examine if total visits indicated a bias for either colour in tested bees. Based on the first choice of a bee to either blue or yellow, a constancy index (CI) was calculated for every individual in each experiment type, using the number of transitions between the two colours in its visit sequence. This index quantified the tendency to switch between colours or to remain consistent with one colour (Slaa et al., 1998) and was calculated as:

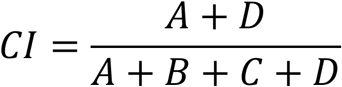

A transition matrix was constructed to categorise transitions such that A and D represent the frequency of transitions within a colour (like transitions: *blue* to *blue* or *yellow* to *yellow*), while B and C represent the frequency of transitions between dissimilar colours (unlike transitions: *blue* to *yellow* or *yellow* to *blue*). CI values range from 0 to 1, where 0 indicates complete inconsistency, 0.5 indicates random foraging and 1 indicates maximum constancy to one colour.

Additionally, we used a Beta regression model to analyse the proportion of total visits to the blue stimulus and the constancy index (CI) across experiments addressing spontaneous constancy under different reward paradigms (*a-c*). The Beta regression was performed using the *betareg* package in R (Cribari-Neto & Zeileis, 2010). The model was specified as the proportion of visits ∼ treatment. Post-hoc and pairwise comparisons were done using the least squares means method with Tukey adjustment by employing the *emmeans* package in R (Searle et al., 1980).

#### Community floral spectra

The proportion of flowers that were binned into short-wavelength and long-wavelength categories were compared using a test of proportions to determine if the community was dominated by one of these categories.

## Results

### Spontaneous constancy to colour

After training to neutral UV-grey associated with low-concentration reward, bees showed a strong bias for blue in tests, with 21 out of 22 bees choosing blue in their first choice (experiment *a*: *χ*^2^ = 16.409, d.f.=1, p<0.001, n=22; Figure 2A) and when total choices were considered (*v* = 231, d.f.=1, p<0.001, n=22; Figure 2B) during tests. Constancy index (CI) was also close to one for bees that chose blue as well as for the single bee that chose yellow (Fig 2C).

**Figure 2.**
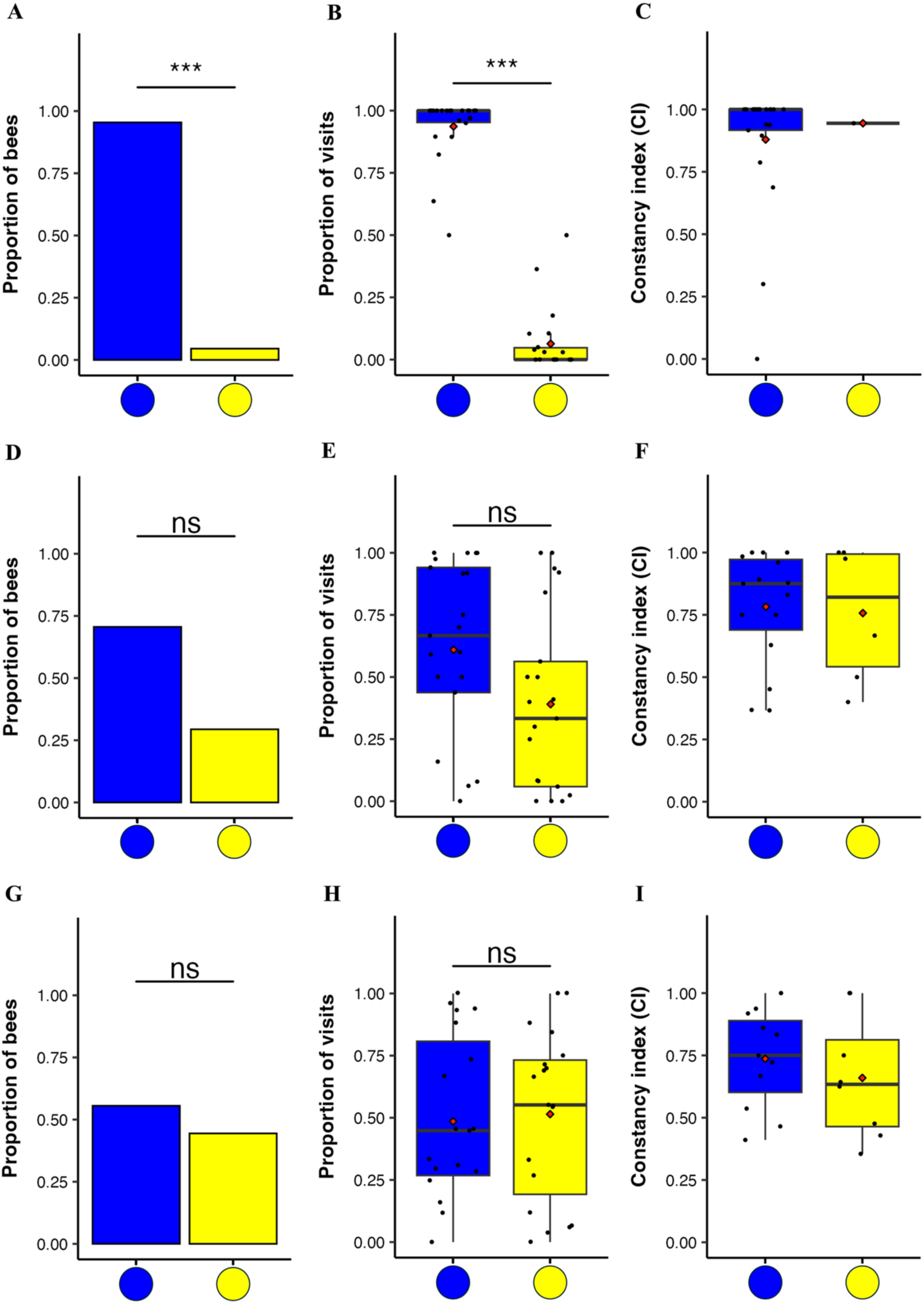
Training was to a UV-grey stimulus in these experiments. *When trained and tested with low concentration reward (experiment a):* A) first choices were strongly biased to blue B) proportion of total visits to either colour was also biased to blue, and the C) constancy of bees based on their first choices for blue or yellow was similar and high. *When trained to high concentration reward and tested with low concentration reward (experiment b):* D) first choices were not significantly biased to blue or yellow, E) total choices were not significantly biased to either colour, and the F) Constancy was high. *When trained and tested with high concentration reward (experiment c):* G) first choices were not significantly biased to either colour, H) Total choices were also not significantly biased to either blue or yellow, and the I) constancy was high. The *hinges* (horizontal bounds of the box) correspond to the interquartile range (IQR), the *bold horizontal line* corresponds to the median and the *whiskers* enclose the range of the data. The black points (•) represent data points for corresponding metrics calculated for individual bees in the trials and the red point (◆) represents the mean. ***depicts significance at p<0.001.

Bees that were trained to UV-grey associated high-concentration reward and tested with low-concentration reward did not show a significant bias for any colour in their first (experiment *b*: *χ*^2^ = 3.048, d.f.=1, p=0.08, n=21; Figure 2D) or total choices (*v* = 121, d.f.=1, p=0.3, n=21; Figure 2E). However, through not statistically significant, an overall preference for blue was observed. Constancy index (CI) was similar and high in bees that chose either colour as their first choice (Mean ± s.d. CI =0.78 ± 0.23, n=21; Figure 2F).

When UV-grey trained bees were both trained and tested with high-concentration reward they did not show a significant bias for either colour in their first (experiment *c*: *χ*^2^ = 0.21, d.f.=1, p=0.64, n=19; Figure 2G) or total choices (*v* = 88, d.f.=1, p=0.79, n=19; Figure 2H). Bees demonstrated high constancy irrespective of whether they chose blue or yellow in their first colour choices (Mean ± s.d.CI =0.7 ± 0.22, n=19; Figure 2I).

### Effect of recent foraging experience on constancy

When bees were trained to yellow, an unpreferred colour compared to the spontaneously preferred blue, their first choices were only to yellow (experiment *d*: *χ*^2^ = 17.05, d.f.=1, p<0.001, n=19; Figure 3A) and total choices were significantly biased to yellow (*v* = 0, d.f.=1, p<0.001, n=19; Figure 3C). Bees also demonstrated a high constancy to yellow (Mean ± s.d.CI = 0.94 ± 0.12, n=19; Figure 3B).

**Figure 3.**
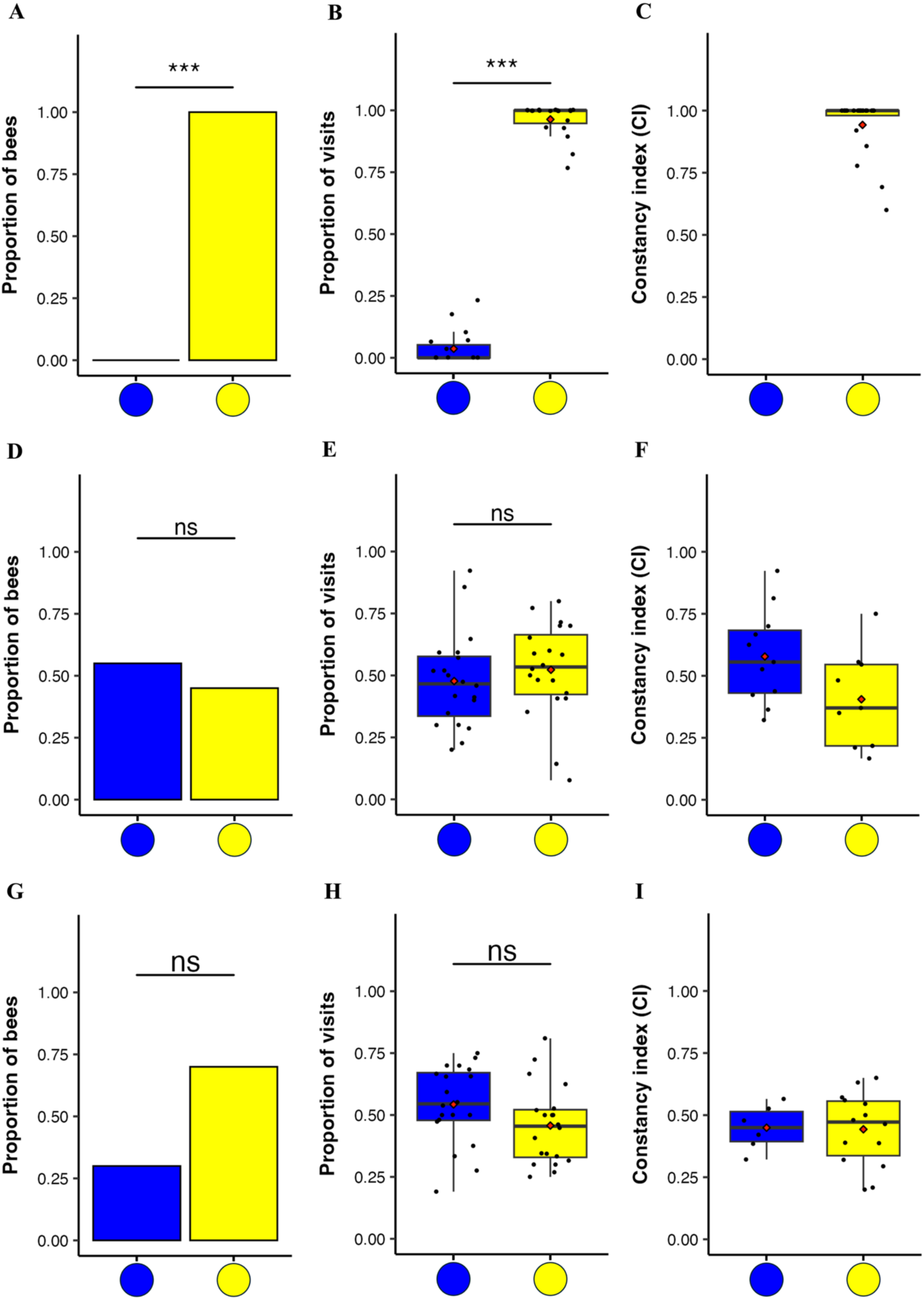
*When trained with yellow colour (experiment d)*: A) first choices were strongly biased to yellow, B) total choices were also biased to yellow, and the C) constancy was high. *When trained to both colours sequentially, with last training to blue*: D) first choices were not biased to blue or yellow, E) total choices were also not biased to blue or yellow, and the F) Constancy was low. *When trained to both colours sequentially, with last training to yellow (experiment e)*: G) first choices were not biased to blue or yellow, H) total choices were also not biased to blue or yellow, and the I) Constancy was low. The *hinges* (horizontal bounds of the box) correspond to the interquartile range (IQR), the *bold horizontal line* corresponds to the median and the *whiskers* enclose the range of the data. The black points (•) represent data points for corresponding metrics calculated for individual bees in the trials and the red point (◆) represents the mean. ***depicts significance at p<0.001.

When trained to both blue and yellow in an alternating sequence, both groups of bees showed no bias for either colour irrespective of whether they were last trained to blue (experiment *e*: *χ*^2^ = 0.05, d.f.=1, p=0.82, n=20; Figure 3D) or yellow (*χ*^2^ = 2.45, d.f.=1, p=0.117, n=20; Figure 3G) in their first choices, as well as in total choices when last trained to blue (*v* = 113, d.f.=1, p=0.47, n=20; Figure 3F) or yellow (*v* = 97.5, d.f.=1, p=0.32, n=20; Figure 3I). Constancy was lower than in the previous experiments in bees that were last trained to blue (Mean ± s.d.CI = 0.37 ± 0.15, n=20; Figure 3E) or yellow (Mean ± s.d.CI = 0.31 ± 0.11, n=20; Figure 3H) suggesting a random foraging pattern.

### Comparison of bee preferences and constancy across experiments

Using beta regression model, we examined the proportion of total visits to the blue stimuli and constancy across experiments (*a-c*). We tested whether there were significant differences in constancy and colour preference between bees trained to UV-grey under various reward paradigms. Preference to blue was significantly reduced when the bees experienced higher quality reward in training alone or both training and test (Table T1: beta regression (formula = proportion of visits ∼ treatment) and pairwise comparison of marginal means with Tukey adjustment, p<0.001). (Supplementary figure 4 (A); Supplementary table 2).

Constancy was high (>0.7) in all three experiments (a-c) (Tables T3-T4; beta regression (formula = constancy index ∼ treatment) and pairwise comparison of marginal means with Tukey adjustment, p<0.001); Supplementary figure 4 (B); Supplementary table 3).

### Community flower spectra

When the floral reflectance spectra was modelled in the bee hexagonal colour space with a green leaf background (Figure 4A), the community spectra was found to be significantly biased towards short wavelengths regardless of whether we considered all flowers (Test of proportions, *χ*^2^ = 18.732, d.f. = 1, p <0.001; Figure 4B), only native species (Test of proportions, *χ*^2^ = 13.846, d.f. = 1, p <0.001; Supplementary figure 5 (A and B)) or only exotic species (Test of proportions, *χ*^2^ = 4.983, d.f. = 1, p = 0.026; Supplementary figure 5 (C and D)).

**Figure 4.**
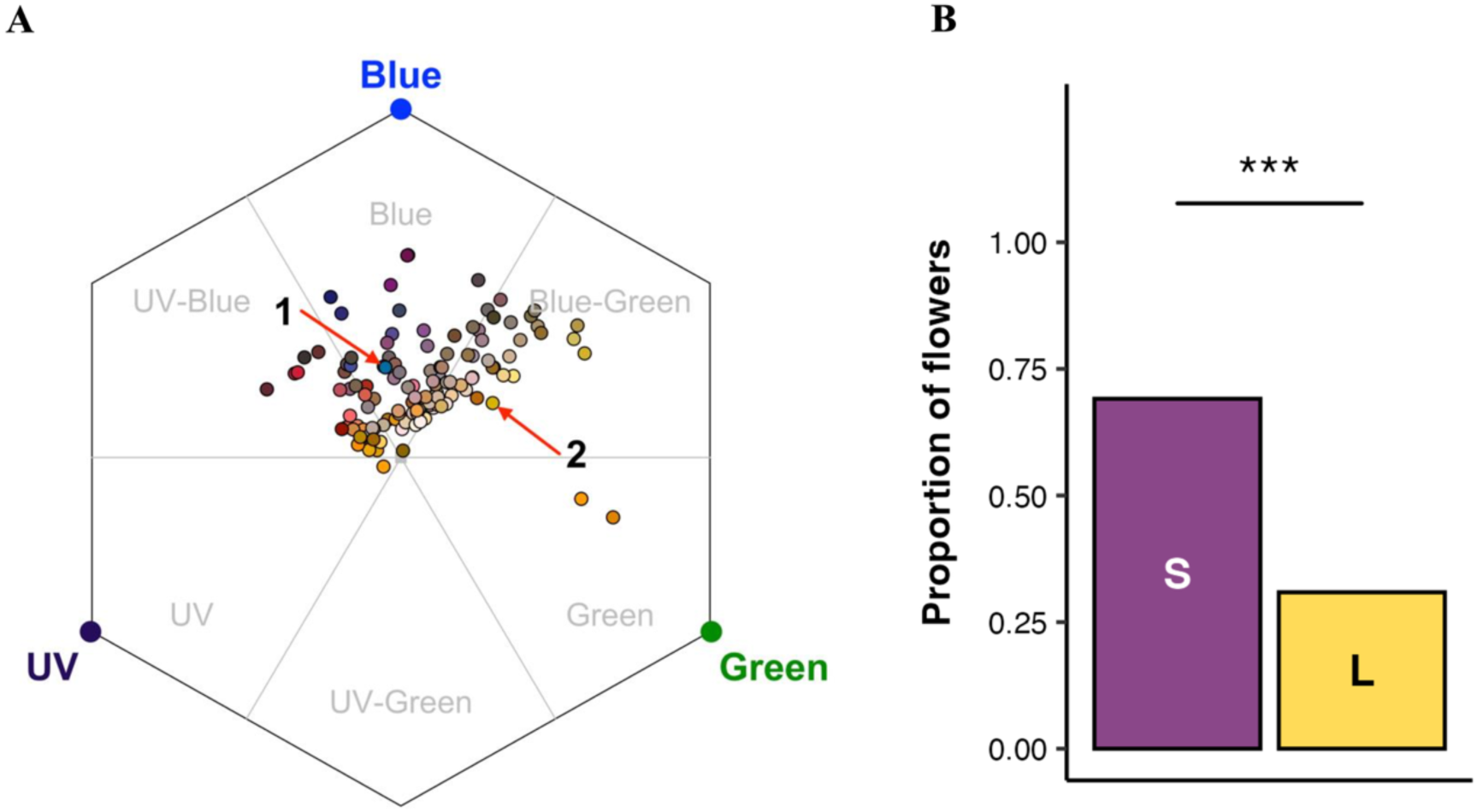
Community flower spectra in the bee subjective hexagonal colour space and categorised into UV, UV-blue, blue, blue-green, green and UV-green sectors: A) A total of 123 flowers in the community were modelled in the bee hexagonal colour space with the background being leaf green. The red arrows point to the blue (labelled 1) and yellow (labelled 2) stimuli used in the experiments. B) flowers categorised into long or short wavelength (labelled “S” and “L” inside the bars) based on the hexagon sector in which the colour loci of the flowers are located. The black points (•) represent the colour loci of the major colour of flowers in the community in the hexagonal colour space. *** depicts significance at p < 0.001.

## Discussion

We examined spontaneous constancy to colour and its modulation by reward concentration and learning in the giant Asian honeybee, *A. dorsata.* Our results highlight an influence of reward concentration on spontaneous colour biases and constancy, which has not been explicitly shown before. When trained to a neutral UV-grey stimulus, bees were strongly biased towards blue and showed high constancy when provided with low-concentration sugar reward during both training and testing. An explanation for this pattern is that, under low reward conditions and in the absence of strong learned associations, bees rely more heavily on their spontaneous colour biases. In our experiments, stimuli were not refilled as a bout progressed, yet bees continued to search on the blue stimuli even after they were depleted, indicating a robust preference for blue which is independent of reward availability. We further found that the community floral spectra were significantly biased towards short-wavelength flowers, suggesting that spontaneous preference for blue may reflect the local floral environment. Interestingly, we found that spontaneous preference for blue was modulated by concentration of the sugar reward bees experienced in our experiments.

Some species of honeybees, bumblebees, stingless bees and butterflies exhibit innate preferences for short-wavelength colours ((Arikawa et al., 2021; Giurfa et al., 1995; Gumbert, 2000; Morawetz et al., 2013; Nicholls et al., 2015; Raine & Chittka, 2005a; Yoshida et al., 2015); but also see (Balamurali et al., 2018)). Bumblebees, for example, are known to revert to their innate colour preference when trained to a neutral stimulus (Giurfa et al., 1995; Gumbert, 2000). Our results from this tropical native honeybee supports this finding. However, we additionally discovered that the strength of the spontaneous blue bias in *A. dorsata* varies significantly with reward concentration; the bias was strongest under low reward concentration and weakened substantially when bees were provided with higher-concentration sugar reward. When bees received brief training on one or both colours (blue or yellow), they subsequently preferred the trained colour in the test, demonstrating that spontaneous preferences can be rapidly overwritten by learning. Modulation of colour preferences in the presence of olfactory cues have been reported in many butterfly species (Anaswara et al., 2025; Yoshida et al., 2015) and multi-sensory integration helps bumblebees locate flowers faster (Kunze & Gumbert, 2001). However, modulation of spontaneous preference by reward quality has not been addressed before. Since reward processing is tightly linked to neuro-modulatory circuits involving biogenic amines in insect brains (Perry & Barron, 2013) it is highly likely that the spontaneous preferences are flexibly controlled by reward quality in bees.

When trained to the neutral UV-grey, bees showed a strong spontaneous preference for blue when they experienced low-concentration reward during training and testing. This bias diminished when bees were trained to a neutral UV-grey paired with high-concentration sugar reward. In previous studies on the Western honeybee *A. mellifera* and the Eastern honeybee *A. cerana* that used similarly high concentrations of sugar solution, typically 30% (w/v or w/w), responses were different from *A. dorsata* in our experiment. *A. mellifera* trained and tested with one molar sugar solution (34.23% w/v or 30.37% w/w) exhibited individual-level constancy, where each individual shows a strong preference for a particular colour, but as a group they exhibited no colour bias (Grüter & Ratnieks, 2011; Wells et al., 1992). In fact, in *A. mellifera* a decrease in reward concentration from 1 to 0.5 molar resulted in reduced constancy but still without a preference for either blue or yellow colour (Grüter et al., 2011). On the other hand, *A. cerana* foraged on both colours and displayed low constancy (Wells & Rathore, 1994). Reward expectation is known to alter foraging preferences, persistence, and exploratory behaviour when bees are presented with unfamiliar stimuli (Gil, 2010; Gil et al., 2007; Solvi et al., 2016). We propose that the increase in reward expectation induced by increasing the reward concentration from 10% w/w to 30% w/w reduced the bees’ reliance on spontaneous colour preferences when confronted with unfamiliar stimuli, prompting them to sample non-preferred colours more frequently. This reward-dependent modulation of spontaneous colour biases has not been reported before.

As expected, spontaneous preference for blue was flexible and readily modified by learning. When trained to the less preferred yellow, bees shifted their preference to the trained yellow in the first and total visits and exhibited high constancy. A brief experience with colour(s) is known to be sufficient for bees to learn it, especially those that contrast sufficiently with the background (Balamurali et al., 2018; Menzel, 1967; Niggebrugge et al., 2009). In our experiments, bees were also able to learn two colours simultaneously during training and used this information when making foraging choices in tests. This is in agreement with findings from *A. mellifera* (Muth et al., 2015, 2017). Previous experiments in similar arrays have attempted to train bees to two colours but have used longer temporal gaps between the training colours which may have biased bees to the last trained colour (Hill et al., 1997). In contrast we show that when training is done to both colours sequentially and in quick succession, bees visit both colour stimuli extensively and also frequently switch between them. This produced a random foraging pattern and correspondingly low constancy, and bees showed no detectable effect of the last trained colour on either the first or total visits. Constraints on the memory of bees, especially short-term memory, have been proposed as a mechanism of constancy (Chittka et al., 1999; Gegear & Laverty, 2001, 2005; Hill et al., 1997, 2001; Waser, 1986). Our results, along with recent studies showing that bees can flexibly learn and use multiple floral cues even over short timescales (Ings et al., 2009; Muth et al., 2015; Raine & Chittka, 2007a), suggest that memory constraints alone may not account for patterns of constancy, at least when floral traits are simple (as in the stimuli used in our experiments).

The bee-subjective floral colours of the community were strongly biased towards short-wavelength hues. This pattern was consistent whether native and exotic species were analysed separately or combined, supporting the idea that the blue preference observed in our experiments is likely innate/spontaneous. Community-wide floral colour assemblages are known to be shaped by the dominant pollinators in a habitat (Ishii et al., 2019; Lunau & Maier, 1995; Reverté et al., 2016), and the colours of the most rewarding flowers often reflect the innate preferences of these naïve pollinators (Chittka et al., 2004; Raine & Chittka, 2007b; Streinzer et al., 2021). The prevalence of flowers in the blue, UV-blue and blue-green sectors in our community is consistent with patterns documented in other regions, where such distributions indicate pollination systems dominated by insects, especially bees (Arnold et al., 2009; Chittka et al., 1994; de Camargo et al., 2019; Dyer et al., 2012, 2021; Ortiz et al., 2021; Shrestha, Burd, et al., 2019; Shrestha, Dyer, et al., 2019; Shrestha et al., 2014).

Innate preferences help naïve pollinators locate flowers and were long assumed to have evolved to guide pollinators toward the most rewarding floral types (Giurfa et al., 1995; Lunau & Maier, 1995; Raine & Chittka, 2007b), although not always (Shrestha et al., 2020). Alternative explanations include mechanisms such as resource partitioning (Balamurali et al., 2018). Short-wavelength preferences have also been proposed to enhance detection of flowers against complex long-wavelength backgrounds (Bukovac et al., 2017), and some studies show that blue flowers are detected and foraged on more efficiently than other colour morphs of the same species (Dyer et al., 2007; Waser & Price, 1985). Taken together, the predominance of blue, UV-blue and blue-green flowers in the community might reflect pollinator-mediated selection on floral colour (Finnell & Koski, 2021; van der Kooi et al., 2016).

It cannot be ruled out that the spontaneous preference for blue in our experiments partly reflects long-term experience with a flower community dominated by short-wavelength colours. Testing truly innate preferences in *A. dorsata* is extremely challenging because, as an open-nesting wild species, obtaining colour-naïve foragers is nearly impossible (Gumbert, 2000) and the experimental methods used may alter these biases (Balamurali et al., 2018). We minimised short-term experience by extensively training bees on neutral UV-grey stimuli (four bouts with 4–16 rewarded landings) and by keeping the intervals between training bouts and testing under two minutes. Both spontaneous and learned preferences shape pollinator foraging (Kinoshita et al., 2017; Raine & Chittka, 2005a). Although spontaneous preferences can initially lead to less efficient choices (such as when the preferred colour is less rewarding or distracts bees from attending to other cues), these effects are unlikely to impose major fitness costs because bees rapidly learn alternative colours, as we and other studies have shown (Menzel, 1985; Morawetz et al., 2013; Muth et al., 2015; Srinivasan, 2010). Thus, while spontaneous preferences can influence constancy and potentially temporarily reduce foraging efficiency, substantial losses are unlikely. Moreover, constancy is influenced by a suite of additional factors, including floral odour, shape, complexity, energetic constraints on learning and memory, social information, and the spatial distribution of flowers (Abts & Dunlap, 2022; Baracchi et al., 2018; Bruninga-Socolar et al., 2022; Gegear & Laverty, 2005; Takagi & Ohashi, 2025). Therefore, constancy can remain an effective and adaptive strategy despite the presence of spontaneous biases.

The artificial flowers used in our experiments were deliberately simple and differed only in colour. This design allows for direct comparison across studies using similar paradigms (Gegear & Laverty, 2004; Grant, 1950; Grüter et al., 2011; Grüter & Ratnieks, 2011; Hill et al., 1997, 2001; Wells et al., 1992; Wells & Rathore, 1994; Wells & Wells, 1983). However, bee behaviour and constancy can shift when flowers vary in morphological complexity, number of morphs, or spatial arrangement (Takagi & Ohashi, 2025). Bees also integrate information from multiple modalities which include colour, odour, shape and size, when they make foraging decisions (for eg., (Gegear & Laverty, 2005; Slaa et al., 1998, 2003)). Future work incorporating multiple morphs and complex multimodal floral signals will be necessary to understand the role of constancy in natural settings.

As is evident from our experiments, *A. dorsata* behaves quite differently even when compared with the closely related temperate honeybee *A. mellifera* and the sympatric tropical honeybee *A. cerana* under similar test conditions (Wells & Rathore, 1994). Such interspecific differences, even among closely related taxa, have been documented previously (Raine & Chittka, 2005b; Slaa et al., 1998; Wells & Rathore, 1994). These findings underscore the importance of species-specific studies for understanding the diversity of foraging behaviours exhibited by different pollinators. Such behavioural variation can have significant consequences for how plant-pollinator interactions are structured across ecological communities.

## Acknowledgements

We thank Balamurali for help with experiments, discussions, and comments on the manuscript. The study was funded by the Anusandhan National Research Foundation (ANRF/SERB SPR/2021/000510) and IISER TVM to HS, and doctoral fellowships from UGC and IISER TVM to SR and SB, respectively.

## Supplementary material

### Figures

**Supplementary figure 1:**
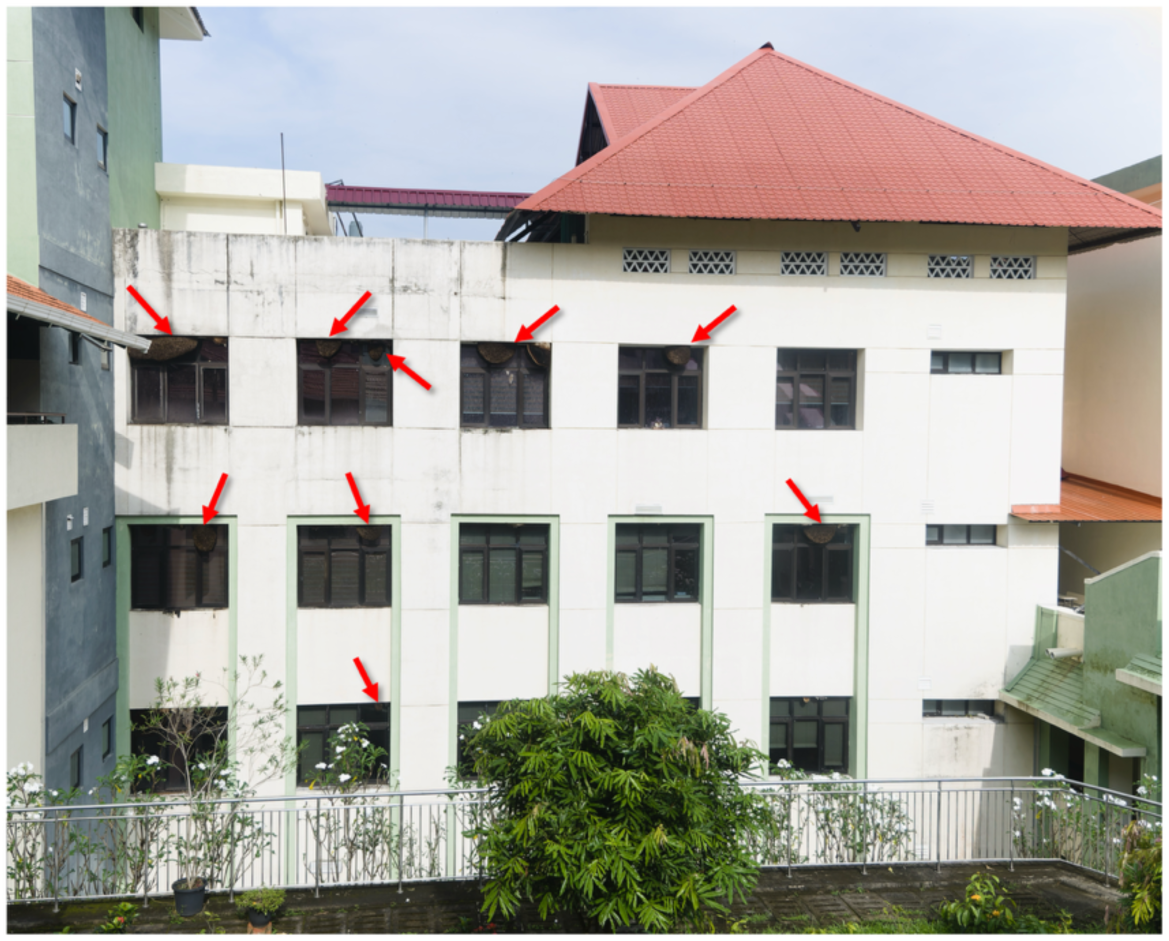
*A. dorsata* colonies on the wall of the biological sciences building at IISER Thiruvananthapuram. The red arrows point to individual colonies from which bees were recruited for experiments.

**Supplementary figure 2:**
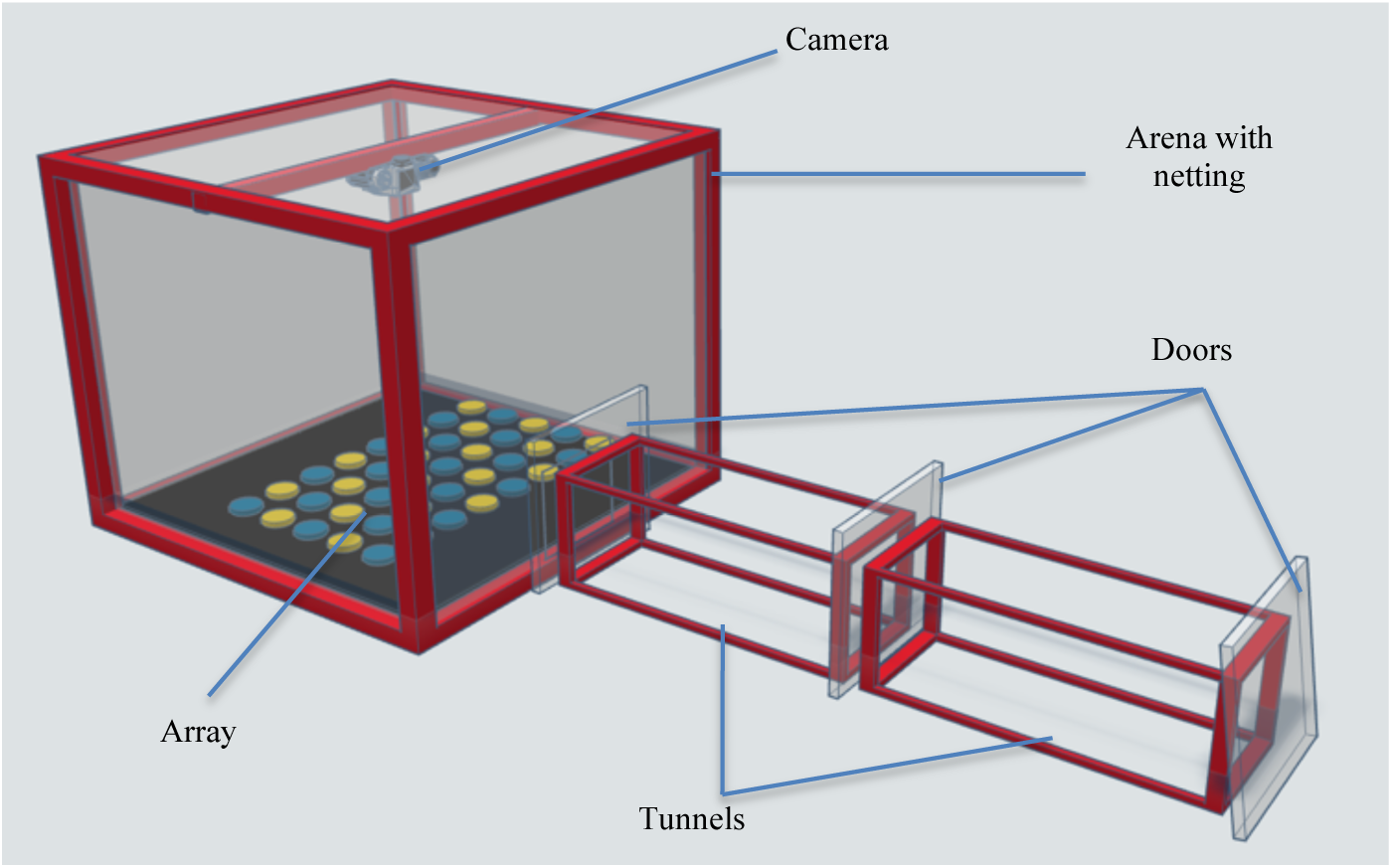
Experimental arena with netting on the walls and a clear plexiglass tunnel leading to the interior of the arena. The three doors in the tunnel help to control the number of bees entering. A camera on the roof is used to record the choices of bees in tests.

**Supplementary figure 3:**
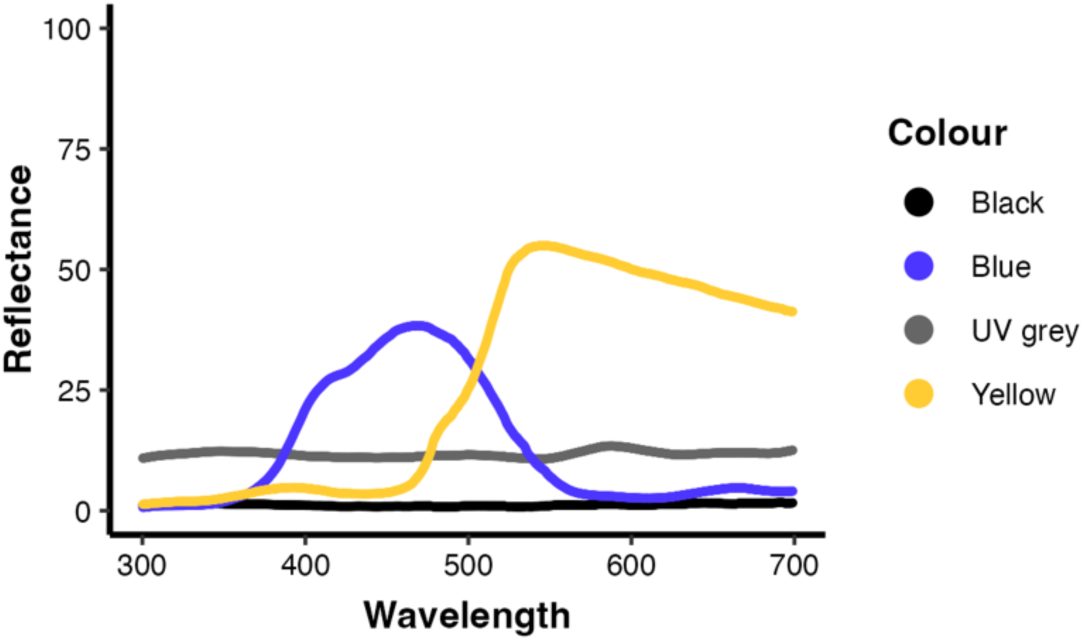
Spectral reflectance of the stimuli and background used in the different experiments

**Supplementary figure 4:**
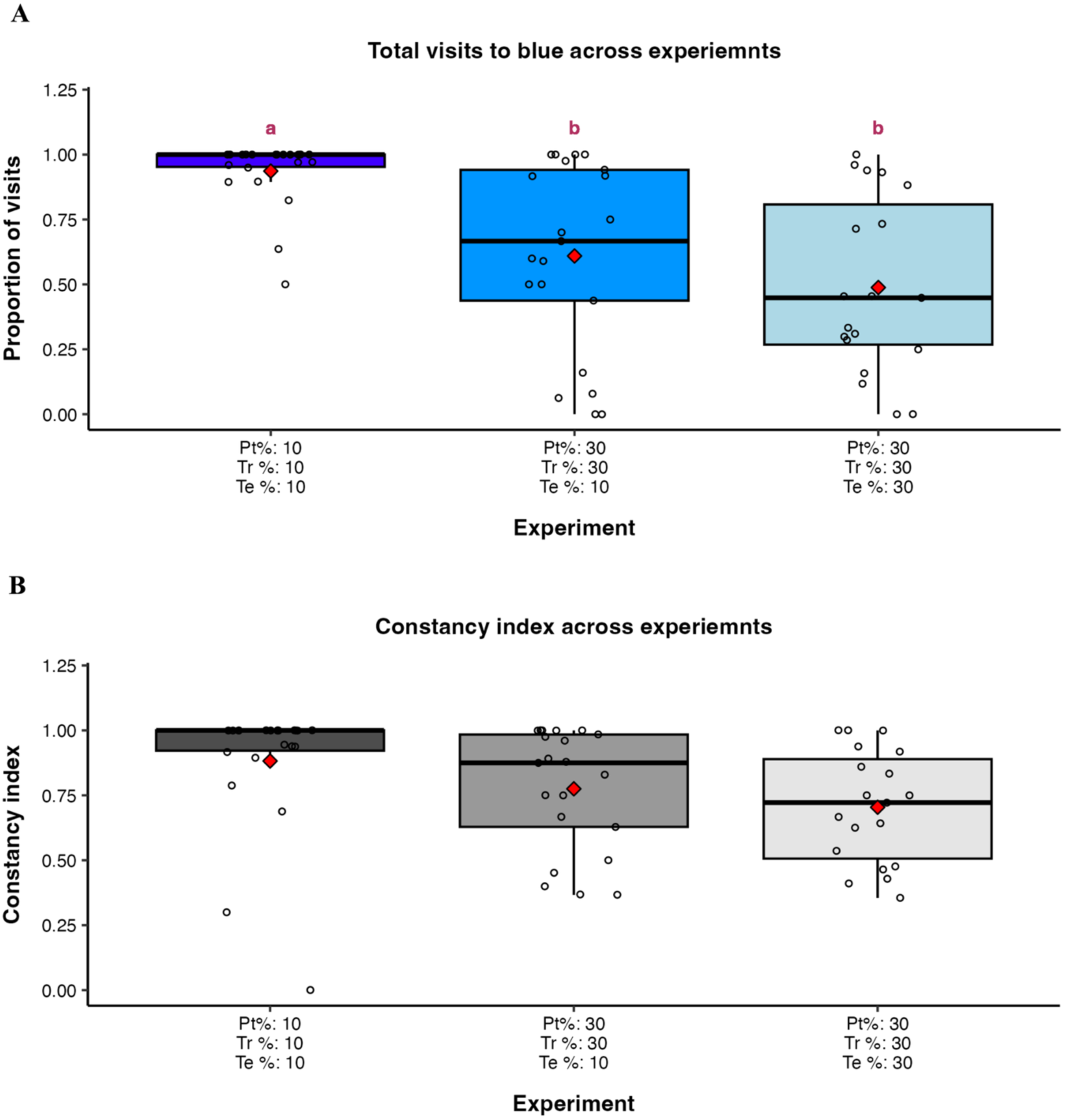
A) Comparison of proportion of visits to blue stimuli in tests across experiments A.1, A.2 and A.3 B) Constancy index comparison across experiments A.1, A.2 and A.3. Pt% stands for pre-training reward concentration, Tr% stands for training reward concentration and Te% stands for test reward concentration. The *hinges* (horizontal bounds of the box) correspond to the interquartile range (IQR), the *bold horizontal line* corresponds to the median and the *whiskers* enclose the range of the data. The open circles (O) represent data points for corresponding metrics calculated for individual bees in the trials and the red point (◆) represents the mean. The letters above the box plots are compact letter display with means not sharing any letter being significantly different using pairwise comparisons employing least squares means method with Tukey adjustment at 5% level of significance.

**Supplementary figure 5:**
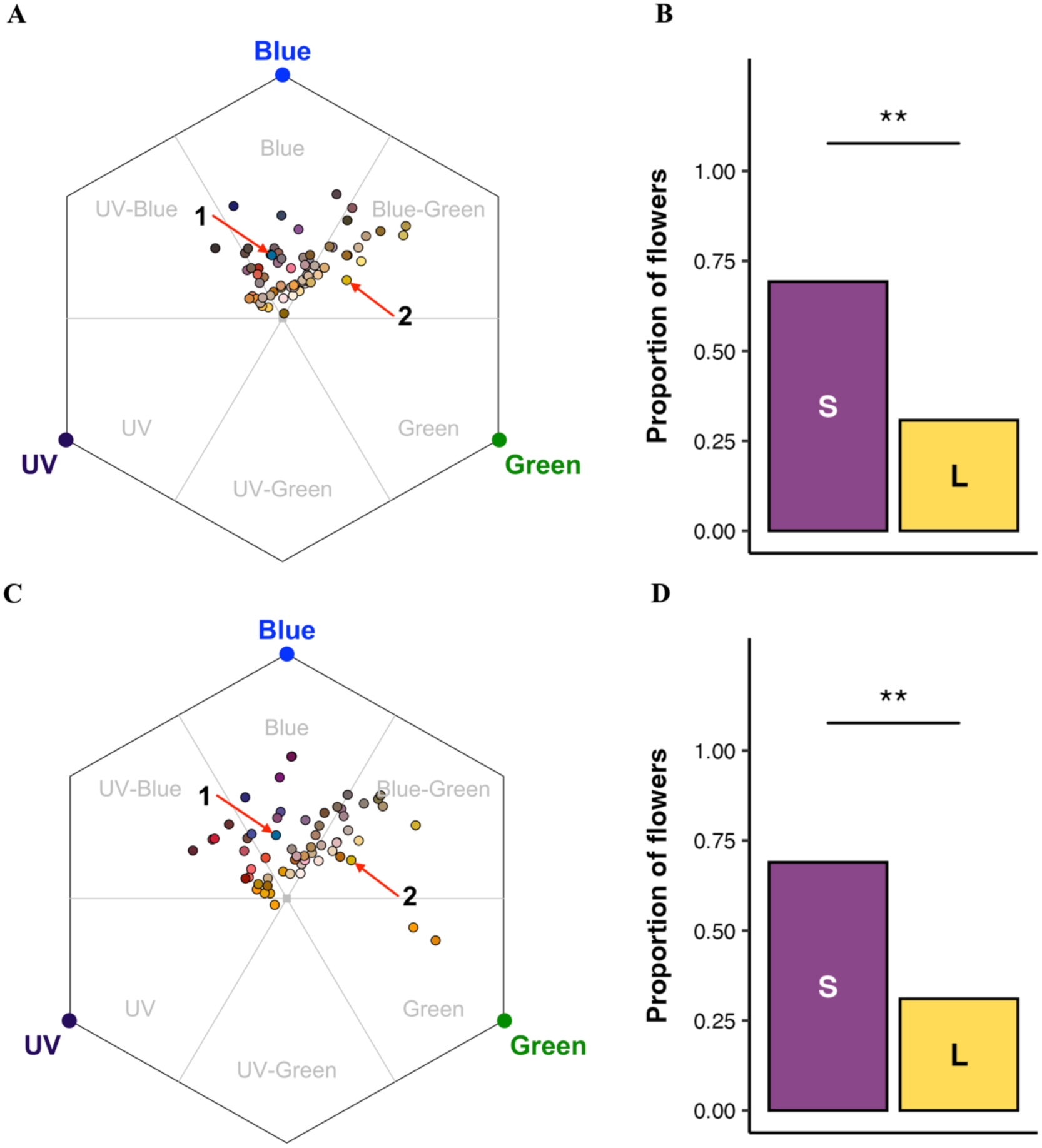
Community flower spectra in the bee subjective hexagonal colour space and categorised into long and short wavelength bee subjective colours A) Native flowers (n=65) modelled in the bee hexagonal colour space with the background being a leaf green. B) The Native flowers (n=65) categorised into short and long wavelength colours (labelled “S” and “L” inside the bars) based on the hexagon sector. C) Exotic flowers (n=58) modelled in the bee hexagonal colour space with the background being a leaf green. D) The Exotic flowers (n=58) categorised into short and long wavelength colours (labelled “S” and “L” inside the bars) based on the hexagon sector. Flowers that fell into UV, UV-blue and blue sectors were classified as short wavelength flowers and those that fell into blue-green, green and UV-green were classified as long wavelength colours. The red arrows point to the blue (labelled 1) and yellow (labelled 2) stimuli used in the experiments. A significant proportion of flowers were represented in the short wavelength category. The black points (•) represent the colour loci of the major colour of flowers in the community in the hexagonal colour space. ** depicts significance at *p* < 0.01.

### Tables

**Table T1:** Colour contrast of the stimuli used in the experiment modelled with black background in the bee-specific Hexagonal colour space.

| Stimuli |  | x | y | r.vec |
| --- | --- | --- | --- | --- |
| 1 | Blue | 0.103 | 0.070 | 0.125 |
| 2 | Yellow | 0.273 | 0.025 | 0.274 |
| 3 | UV-grey | 0.018 | -0.001 | 0.018 |

**Table T2:** Pairwise comparison of proportion of total visits to the blue colour stimuli across experiments a,b and c with beta regression (formula = frequency of visits ∼ treatment) using marginal means method and Tukey adjustment.

**Quantile residuals:**
| Min | 1Q | Median | 3Q | Max |
| --- | --- | --- | --- | --- |
| -2.455 | -0.340 | -0.015 | 0.603 | 2.149 |

**Coefficients:**
| component |  | term | estimate | SE | statistic | p value |
| --- | --- | --- | --- | --- | --- | --- |
| 1 | mean | *Experiment a* (Intercept) | 1.542 | 0.274 | 5.620 | 1.91e-08 |
| 2 | mean | *Experiment b*<br>Training concentration: 30%<br>Test concentration: 10% | -1.129 | 0.376 | -3.000 | 0.003 |
| 3 | mean | *Experiment c*<br>Training concentration: 30%<br>Test concentration: 30% | -1.609 | 0.402 | -4.001 | 6.31e-05 |

**Emmeans:**
|  | experiment | estimate | SE | 95% CI<br>(Lower) | 95% CI<br>(Upper) |
| --- | --- | --- | --- | --- | --- |
|  | *Experiment a* |  |  |  |  |
| 1 | Training concentration: 10%<br>Test concentration: 10% | 0.824 | 0.040 | 0.746 | 0.902 |
|  | *Experiment b* |  |  |  |  |
| 2 | Training concentration: 30%<br>Test concentration: 10% | 0.602 | 0.066 | 0.472 | 0.732 |
|  | *Experiment c* |  |  |  |  |
| 3 | Training concentration: 30%<br>Test concentration: 30% | 0.483 | 0.073 | 0.341 | 0.626 |

**Pairwise contrast:**
|  | contrast | estimate | SE | Z-ratio | p value |
| --- | --- | --- | --- | --- | --- |
|  | *Experiment a* |  |  |  |  |
|  | Training concentration: 10%<br>Test concentration: 10% |  |  |  |  |
| 1 | vs | 0.222 | 0.075 | 2.955 | 0.009 |
|  | *Experiment b* |  |  |  |  |
|  | Training concentration: 30%<br>Test concentration: 10% |  |  |  |  |
|  | *Experiment a* |  |  |  |  |
|  | Training concentration: 10%<br>Test concentration: 10% |  |  |  |  |
| 2 | vs | 0.340 | 0.083 | 4.090 | 0.000 |
|  | *Experiment c* |  |  |  |  |
|  | Training concentration: 30%<br>Test concentration: 30% |  |  |  |  |
|  | *Experiment b* |  |  |  |  |
|  | Training concentration: 30%<br>Test concentration: 10% |  |  |  |  |
| 3 | vs | 0.119 | 0.099 | 1.201 | 0.453 |
|  | *Experiment c* |  |  |  |  |
|  | Training concentration: 30%<br>Test concentration: 30% |  |  |  |  |

**Table T3:** Pairwise comparison of constancy index across experiments a, b and c with beta regression (formula = CI ∼ treatment) using marginal means and Tukey adjustment.

**Quantile residuals:**
| Min | 1Q | Median | 3Q | Max |
| --- | --- | --- | --- | --- |
| -4.062 | -0.589 | -0.078 | 0.772 | 1.703 |

**Coefficients:**
| component |  | term | estimate | SE | statistic | p value |
| --- | --- | --- | --- | --- | --- | --- |
| 1 | mean | *Experiment a* (Intercept) | 1.727 | 0.272 | 6.344 | 2.24e- |
|  |  | *Experiment b* |  |  |  |  |
| 2 | mean | Training concentration: 30% |  |  |  |  |
|  |  | Test concentration: 10% | -0.527 | 0.345 | -1.527 | 0.127 |
|  |  | *Experiment c* |  |  |  |  |
| 3 | mean | Training concentration: 30% |  |  |  |  |
|  |  | Test concentration: 30% | -0.872 | 0.360 | -2.422 | 0.015 |

**Emmeans:**
|  | experiment | estimate | SE | 95% CI<br>(Lower) | 95% CI<br>(Upper) |
| --- | --- | --- | --- | --- | --- |
|  | *Experiment a* |  |  |  |  |
| 1 | Training concentration: 10%<br>Test concentration: 10% | 0.849 | 0.035 | 0.781 | 0.917 |
|  | *Experiment b* |  |  |  |  |
| 2 | Training concentration: 30%<br>Test concentration: 10% | 0.769 | 0.048 | 0.674 | 0.863 |
|  | *Experiment c* |  |  |  |  |
| 3 | Training concentration: 30%<br>Test concentration: 30% | 0.702 | 0.058 | 0.588 | 0.815 |

**Pairwise contrast:**
|  | contrast | estimate | SE | Z-ratio | p value |
| --- | --- | --- | --- | --- | --- |
|  | *Experiment a* |  |  |  |  |
|  | Training concentration: 10%<br>Test concentration: 10% |  |  |  |  |
| 1 | vs | 0.080 | 0.054 | 1.498 | 0.292 |
|  | *Experiment b* |  |  |  |  |
|  | Training concentration: 30%<br>Test concentration: 10% |  |  |  |  |
|  | *Experiment a* |  |  |  |  |
|  | Training concentration: 10%<br>Test concentration: 10% |  |  |  |  |
| 2 | vs | 0.147 | 0.063 | 2.327 | 0.052 |
|  | *Experiment c* |  |  |  |  |
|  | Training concentration: 30%<br>Test concentration: 30% |  |  |  |  |
|  | *Experiment b* |  |  |  |  |
|  | Training concentration: 30%<br>Test concentration: 10% |  |  |  |  |
| 3 | vs | 0.067 | 0.071 | 0.942 | 0.614 |
|  | *Experiment c* |  |  |  |  |
|  | Training concentration: 30%<br>Test concentration: 30% |  |  |  |  |

## References

1. Abts, B. J., & Dunlap, A. S. (2022). Memory and the value of social information in foraging bumble bees. Learning & Behavior, 50(3), 317–328. 10.3758/s13420-022-00528-2

2. Amaya-Márquez, M. (2009). Floral constancy in bees: A revision of theories and a comparison with other pollinators. 1.

3. Anaswara, K., Balamurali, G., Somanathan, H., & Kodandaramaiah, U. (2025). Reproductive status modulates colour preference and multimodal cue integration in host plant location by butterflies. Journal of Experimental Biology, 228(22), jeb250414.

4. Arikawa, K., Nakatani, Y., Koshitaka, H., & Kinoshita, M. (2021). Foraging Small White Butterflies, *Pieris rapae*, Search Flowers Using Color Vision. Frontiers in Ecology and Evolution, 9, 650069. 10.3389/fevo.2021.650069

5. Arnold, S. E., Savolainen, V., & Chittka, L. (2009). Flower colours along an alpine altitude gradient, seen through the eyes of fly and bee pollinators. Arthropod-Plant Interactions, 3, 27–43.

6. Balamurali, G. S., Nicholls, E., Somanathan, H., & Hempel De Ibarra, N. (2018). A comparative analysis of colour preferences in temperate and tropical social bees. The Science of Nature, 105(1–2), 8. 10.1007/s00114-017-1531-z

7. Baracchi, D., Vasas, V., Jamshed Iqbal, S., & Alem, S. (2018). Foraging bumblebees use social cues more when the task is difficult. Behavioral Ecology, 29(1), 186–192. 10.1093/beheco/arx143

8. Baude, M., Danchin, É., Mugabo, M., & Dajoz, I. (2011). Conspecifics as informers and competitors: An experimental study in foraging bumble-bees. Proceedings of the Royal Society B: Biological Sciences, 278(1719), 2806–2813. 10.1098/rspb.2010.2659

9. Bruninga-Socolar, B., Winfree, R., & Crone, E. E. (2022). The contribution of plant spatial arrangement to bumble bee flower constancy. Oecologia, 198(2), 471–481.

10. Bukovac, Z., Shrestha, M., Garcia, J. E., Burd, M., Dorin, A., & Dyer, A. G. (2017). Why background colour matters to bees and flowers. Journal of Comparative Physiology A, 203, 369–380.

11. Chittka, L. (1992). The colour hexagon: A chromaticity diagram based on photoreceptor excitations as a generalized representation of colour opponency. Journal of Comparative Physiology A, 170(5). 10.1007/BF00199331

12. Chittka, L., Ings, T. C., & Raine, N. E. (2004). Chance and adaptation in the evolution of island bumblebee behaviour. Population Ecology, 46, 243–251.

13. Chittka, L., & Menzel, R. (1992). The evolutionary adaptation of flower colours and the insect pollinators’ colour vision. Journal of Comparative Physiology A, 171(2). 10.1007/BF00188925

14. Chittka, L., & Raine, N. E. (2006). Recognition of flowers by pollinators. Current Opinion in Plant Biology, 9(4), 428–435.

15. Chittka, L., Shmida, A., Troje, N., & Menzel, R. (1994). Ultraviolet as a component of flower reflections, and the colour perception of Hymenoptera. Vision Research, 34(11), 1489–1508.

16. Chittka, L., Thomson, J. D., & Waser, N. M. (1999). Flower Constancy, Insect Psychology, and Plant Evolution. Naturwissenschaften, 86(8), 361–377. 10.1007/s001140050636

17. Cribari-Neto, F., & Zeileis, A. (2010). Beta regression in R. Journal of Statistical Software, 34, 1–24.

18. de Camargo, M. G. G., Lunau, K., Batalha, M. A., Brings, S., de Brito, V. L. G., & Morellato, L. P. C. (2019). How flower colour signals allure bees and hummingbirds: A community-level test of the bee avoidance hypothesis. New Phytologist, 222(2), 1112–1122.

19. De Jager, M. L., Dreyer, L. L., & Ellis, A. G. (2011). Do pollinators influence the assembly of flower colours within plant communities? Oecologia, 166(2), 543–553.

20. Dyer, A. G., Boyd-Gerny, S., McLoughlin, S., Rosa, M. G., Simonov, V., & Wong, B. B. (2012). Parallel evolution of angiosperm colour signals: Common evolutionary pressures linked to hymenopteran vision. Proceedings of the Royal Society B: Biological Sciences, 279(1742), 3606–3615.

21. Dyer, A. G., & Chittka, L. (2004). Bumblebees (*Bombus terrestris*) sacrifice foraging speed to solve difficult colour discrimination tasks. Journal of Comparative Physiology A, 190(9), 759–763.

22. Dyer, A. G., Jentsch, A., Burd, M., Garcia, J. E., Giejsztowt, J., Camargo, M. G., Tjørve, E., Tjørve, K. M., White, P., & Shrestha, M. (2021). Fragmentary blue: Resolving the rarity paradox in flower colors. Frontiers in Plant Science, 11, 618203.

23. Dyer, A. G., Whitney, H. M., Arnold, S. E., Glover, B. J., & Chittka, L. (2007). Mutations perturbing petal cell shape and anthocyanin synthesis influence bumblebee perception of *Antirrhinum majus* flower colour. Arthropod-Plant Interactions, 1, 45–55.

24. Finnell, L. M., & Koski, M. H. (2021). A test of Sensory Drive in plant–pollinator interactions: Heterogeneity in the signalling environment shapes pollinator preference for a floral visual signal. New Phytologist, 232(3), 1436–1448. 10.1111/nph.17631

25. Gegear, R. J., & Laverty, T. M. (2001). The effect of variation among floral traits on the flower constancy of pollinators. In L. Chittka & J. D. E. Thomson (Eds.), Cognitive ecology of pollination: Animal behaviour and floral evolution (pp. 1–20). Cambridge University Press.

26. Gegear, R. J., & Laverty, T. M. (2004). Effect of a colour dimorphism on the flower constancy of honey bees and bumble bees. Canadian Journal of Zoology, 82(4), 587–593.

27. Gegear, R. J., & Laverty, T. M. (2005). Flower constancy in bumblebees: A test of the trait variability hypothesis. Animal Behaviour, 69(4), 939–949.

28. Gil, M. (2010). Reward expectations in honeybees. Communicative & Integrative Biology, 3(2), 95–100. 10.4161/cib.3.2.10621

29. Gil, M., Marco, R. J. D., & Menzel, R. (2007). Learning reward expectations in honeybees. Learning & Memory, 14(7), 491–496. 10.1101/lm.618907

30. Giurfa, M., Nunez, J., Chittka, L., & Menzel, R. (1995). Colour preferences of flower-naive honeybees. Journal of Comparative Physiology A, 177, 247–259.

31. Goulson, D. (2000). Are insects flower constant because they use search images to find flowers? Oikos, 88(3), 547–552.

32. Goulson, D., & Cory, J. S. (1993). Flower constancy and learning in foraging preferences of the green-veined white butterfly *Pleris napi*. Ecological Entomology, 18(4), 315–320. 10.1111/j.1365-2311.1993.tb01107.x

33. Goulson, D., Stout, J. C., & Hawson, S. A. (1997). Can flower constancy in nectaring butterflies be explained by Darwin’s interference hypothesis? Oecologia, 112(2), 225–231. 10.1007/s004420050304

34. Goulson, D., & Wright, N. P. (1998). Flower constancy in the hoverflies *Episyrphus balteatus* (Degeer) and *Syrphus ribesii* (L.)(Syrphidae). Behavioral Ecology, 9(3), 213–219.

35. Grant, V. (1950). The flower constancy of bees. The Botanical Review, 16(7), 379–398. 10.1007/BF02869992

36. Grüter, C., Moore, H., Firmin, N., Helanterä, H., & Ratnieks, F. L. (2011). Flower constancy in honey bee workers (*Apis mellifera*) depends on ecologically realistic rewards. Journal of Experimental Biology, 214(8), 1397–1402.

37. Grüter, C., & Ratnieks, F. L. W. (2011). Flower constancy in insect pollinators: Adaptive foraging behaviour or cognitive limitation? Communicative & Integrative Biology, 4(6), 633–636. 10.4161/cib.16972

38. Gumbert, A. (2000). Color choices by bumble bees (*Bombus terrestris*): Innate preferences and generalization after learning. Behavioral Ecology and Sociobiology, 48, 36–43.

39. Hayes, L., & Grüter, C. (2023). When should bees be flower constant? An agent-based model highlights the importance of social information and foraging conditions. Journal of Animal Ecology, 92(3), 580–593. 10.1111/1365-2656.13861

40. Heinrich, B., Mudge, P. R., & Deringis, P. G. (1977). Laboratory analysis of flower constancy in foraging bumblebees: *Bombus ternarius* and *B. terricola*. Behavioral Ecology and Sociobiology, 247–265.

41. Hill, P. S. M., Hollis, J., & Wells, H. (2001). Foraging decisions in nectarivores: Unexpected interactions between flower constancy and energetic rewards. Animal Behaviour, 62(4), 729–737. 10.1006/anbe.2001.1775

42. Hill, P. S. M., Wells, P. H., & Wells, H. (1997). Spontaneous flower constancy and learning in honey bees as a function of colour. Animal Behaviour, 54(3), 615–627. 10.1006/anbe.1996.0467

43. Ings, T. C., Raine, N. E., & Chittka, L. (2009). A population comparison of the strength and persistence of innate colour preference and learning speed in the bumblebee *Bombus terrestris*. Behavioral Ecology and Sociobiology, 63(8), 1207–1218.

44. Ishii, H. S. (2005). Analysis of bumblebee visitation sequences within single bouts: Implication of the overstrike effect on short-term memory. Behavioral Ecology and Sociobiology, 57(6), 599–610. 10.1007/s00265-004-0889-z

45. Ishii, H. S., & Kadoya, E. Z. (2016). Legitimate visitors and nectar robbers on *Trifolium pratense* showed contrasting flower fidelity versus co-flowering plant species: Could motor learning be a major determinant of flower constancy by bumble bees? Behavioral Ecology and Sociobiology, 70(3), 377–386. 10.1007/s00265-016-2057-7

46. Ishii, H. S., Kubota, M. X., Tsujimoto, S. G., & Kudo, G. (2019). Association between community assemblage of flower colours and pollinator fauna: A comparison between Japanese and New Zealand alpine plant communities. Annals of Botany, 123(3), 533–541. 10.1093/aob/mcy188

47. Ishii, H. S., & Masuda, H. (2014). Effect of flower visual angle on flower constancy: A test of the search image hypothesis. Behavioral Ecology, 25(4), 933–944. 10.1093/beheco/aru071

48. Kinoshita, M., Stewart, F. J., & Ômura, H. (2017). Multisensory integration in Lepidoptera: Insights into flower-visitor interactions. BioEssays, 39(4), 1600086. 10.1002/bies.201600086

49. Kunze, J., & Gumbert, A. (2001). The combined effect of color and odor on flower choice behavior of bumble bees in flower mimicry systems. Behavioral Ecology, 12(4), 447–456.

50. Lewis, A. C. (1986). Memory constraints and flower choice in *Pieris rapae*. Science, 232(4752), 863–865.

51. Lunau, K., & Maier, E. J. (1995). Innate colour preferences of flower visitors. Journal of Comparative Physiology A, 177(1). 10.1007/BF00243394

52. Maia, R., Eliason, C. M., Bitton, P., Doucet, S. M., & Shawkey, M. D. (2013). pavo: An R package for the analysis, visualization and organization of spectral data. Methods in Ecology and Evolution, 4(10), 906–913. 10.1111/2041-210X.12069

53. Menzel, R. (1967). Untersuchungen zum Erlernen von Spektralfarben durch die Honigbiene (*Apis mellifica*). Zeitschrift f r Vergleichende Physiologie, 56(1), 22–62.

54. Menzel, R. (1985). Learning in honey bees in an ecological and behavioral context. Fortschritte Der Zoologie (Stuttgart).

55. Menzel, R., & Shmida, A. (1993). The ecology of flower colours and the natural colour vision of insect pollinators: The Israeli flora as a study case. Biological Reviews, 68(1), 81–120.

56. Morawetz, L., Svoboda, A., Spaethe, J., & Dyer, A. G. (2013). Blue colour preference in honeybees distracts visual attention for learning closed shapes. Journal of Comparative Physiology A, 199, 817–827.

57. Morellato, L. P. C., Camargo, M. G. G., & Gressler, E. (2013). A review of plant phenology in South and Central America. Phenology: An Integrative Environmental Science, 91–113.

58. Muth, F., Papaj, D. R., & Leonard, A. S. (2015). Colour learning when foraging for nectar and pollen: Bees learn two colours at once. Biology Letters, 11(9), 20150628.

59. Muth, F., Papaj, D. R., & Leonard, A. S. (2017). Multiple rewards have asymmetric effects on learning in bumblebees. Animal Behaviour, 126, 123–133.

60. Newstrom, L. E., Frankie, G. W., & Baker, H. G. (1994). A new classification for plant phenology based on flowering patterns in lowland tropical rain forest trees at La Selva, Costa Rica. Biotropica, 141–159.

61. Nicholls, E. K., Ehrendreich, D., & Hempel De Ibarra, N. (2015). Differences in color learning between pollen- and sucrose-rewarded bees. Communicative & Integrative Biology, 8(4), e1052921. 10.1080/19420889.2015.1052921

62. Niggebrugge, C., Leboulle, G., Menzel, R., Komischke, B., & de Ibarra, N. H. (2009). Fast learning but coarse discrimination of colours in restrained honeybees. Journal of Experimental Biology, 212(9), 1344–1350.

63. Nityananda, V., & Chittka, L. (2021). Different effects of reward value and saliency during bumblebee visual search for multiple rewarding targets. Animal Cognition, 24(4), 803–814.

64. Ogilvie, J. E., & Forrest, J. R. (2017). Interactions between bee foraging and floral resource phenology shape bee populations and communities. Current Opinion in Insect Science, 21, 75–82.

65. Ollerton, J., Johnson, S. D., & Hingston, A. B. (2006). Geographical variation in diversity and specificity of pollination systems. Plant–Pollinator Interactions: From Specialization to Generalization, 283–308.

66. Ortiz, P. L., Fernández-Díaz, P., Pareja, D., Escudero, M., & Arista, M. (2021). Do visual traits honestly signal floral rewards at community level? Functional Ecology, 35(2), 369–383.

67. Perry, C. J., & Barron, A. B. (2013). Neural mechanisms of reward in insects. Annual Review of Entomology, 58(1), 543–562.

68. Plants of the World Online | Kew Science. (2025). Plants of the World Online. https://powo.science.kew.org/

69. R Core Team. (2025). *R: a language and environment for statistical computing* [Manual]. R Foundation for Statistical Computing. https://www.R-project.org/

70. Raine, N. E., & Chittka, L. (2005a). Colour preferences in relation to the foraging performance and fitness of the bumblebee *Bombus terrestris*. Uludağ Arıcılık Dergisi, 5(4), 145–150.

71. Raine, N. E., & Chittka, L. (2005b). Comparison of flower constancy and foraging performance in three bumblebee species (Hymenoptera: Apidae: Bombus). Entomologia Generalis, 28(2), 081.

72. Raine, N. E., & Chittka, L. (2007a). Flower constancy and memory dynamics in bumblebees (Hymenoptera: Apidae: Bombus). Entomol Gen.

73. Raine, N. E., & Chittka, L. (2007b). The adaptive significance of sensory bias in a foraging context: Floral colour preferences in the bumblebee *Bombus terrestris*. PLoS One, 2(6), e556.

74. Reverté, S., Retana, J., Gómez, J. M., & Bosch, J. (2016). Pollinators show flower colour preferences but flowers with similar colours do not attract similar pollinators. Annals of Botany, 118(2), 249–257. 10.1093/aob/mcw103

75. Schmid, B., Nottebrock, H., Esler, K. J., Pagel, J., Böhning-Gaese, K., Schurr, F. M., Mueller, T., & Schleuning, M. (2016). A bird pollinator shows positive frequency dependence and constancy of species choice in natural plant communities. Ecology, 97(11), 3110–3118.

76. Searle, S. R., Speed, F. M., & Milliken, G. A. (1980). Population Marginal Means in the Linear Model: An Alternative to Least Squares Means. The American Statistician, 34(4), 216–221. 10.1080/00031305.1980.10483031

77. Shrestha, M., Burd, M., Garcia, J. E., Dorin, A., & Dyer, A. G. (2019). Colour evolution within orchids depends on whether the pollinator is a bee or a fly. Plant Biology, 21(4), 745–752.

78. Shrestha, M., Dyer, A. G., Bhattarai, P., & Burd, M. (2014). Flower colour and phylogeny along an altitudinal gradient in the Himalayas of Nepal. Journal of Ecology, 102(1), 126–135.

79. Shrestha, M., Dyer, A. G., Garcia, J. E., & Burd, M. (2019). Floral colour structure in two Australian herbaceous communities: It depends on who is looking. Annals of Botany, 124(2), 221–232.

80. Shrestha, M., Garcia, J. E., Burd, M., & Dyer, A. G. (2020). Australian native flower colours: Does nectar reward drive bee pollinator flower preferences? PLOS ONE, 15(6), e0226469. 10.1371/journal.pone.0226469

81. Shrotri, S., Kaur, S., Nawge, V., Sandhya, S., Dandavate, R., & Gowda, V. (2024). Revisiting Aristotle’s observation on bees: High floral constancy is common among bees but it is shaped by the locally abundant flowering species (p. 2024.10.28.614270). bioRxiv. 10.1101/2024.10.28.614270

82. Slaa, Cevaal, A., & Sommeijer, M. J. (1998). Floral constancy in Trigona stingless bees foraging on artificial flower patches: A comparative study. Journal of Apicultural Research, 37(3), 191–198.

83. Slaa, Tack, A. J., & Sommeijer, M. J. (2003). The effect of intrinsic and extrinsic factors on flower constancy in stingless bees. Apidologie, 34(5), 457–468.

84. Solvi, C., Baciadonna, L., & Chittka, L. (2016). Unexpected rewards induce dopamine-dependent positive emotion–like state changes in bumblebees. Science, 353(6307), 1529–1531. 10.1126/science.aaf4454

85. Somanathan, H. (2024). Why diversity matters for understanding the visual ecology and behaviour of bees. Current Opinion in Insect Science, 64, 101224. 10.1016/j.cois.2024.101224

86. Somanathan, H., & Balamurali, G. S. (2023). Comparative Psychophysics of Colour Preferences and Colour Learning in Bees with Special Focus on Asian Social Bees. Journal of the Indian Institute of Science, 103(4), 971–980. 10.1007/s41745-023-00386-5

87. Somanathan, H., Krishna, S., Jos, E. M., Gowda, V., Kelber, A., & Borges, R. M. (2020). Nocturnal bees feed on diurnal leftovers and pay the price of day–night lifestyle transition. Frontiers in Ecology and Evolution, 8, 566964.

88. Somanathan, H., Saryan, P., & Balamurali, G. S. (2019). Foraging strategies and physiological adaptations in large carpenter bees. Journal of Comparative Physiology A, 205(3), 387–398. 10.1007/s00359-019-01323-7

89. Srinivasan, M. V. (2010). Honey Bees as a Model for Vision, Perception, and Cognition. Annual Review of Entomology, 55(Volume 55, 2010), 267–284. 10.1146/annurev.ento.010908.164537

90. Streinzer, M., Neumayer, J., & Spaethe, J. (2021). Flower Color as Predictor for Nectar Reward Quantity in an Alpine Flower Community. Frontiers in Ecology and Evolution, 9, 721241. 10.3389/fevo.2021.721241

91. Takagi, K., & Ohashi, K. (2025). Realized flower constancy in bumble bees: Optimal foraging strategy balancing cognitive and travel costs and its possible consequences for floral diversity. Functional Ecology, 39(3), 863–875. 10.1111/1365-2435.70008

92. Thomson, J. D. (1981). Field measures of flower constancy in bumblebees. American Midland Naturalist, 377–380.

93. van der Kooi, C. J., Dyer, A. G., Kevan, P. G., & Lunau, K. (2019). Functional significance of the optical properties of flowers for visual signalling. Annals of Botany, 123(2), 263–276. 10.1093/aob/mcy119

94. van der Kooi, C. J., Pen, I., Staal, M., Stavenga, D. G., & Elzenga, J. T. M. (2016). Competition for pollinators and intra-communal spectral dissimilarity of flowers. Plant Biology, 18(1), 56–62.

95. van der Niet, T., Pires, K., & Steenhuisen, S.-L. (2020). Flower constancy of the Cape honey bee pollinator of two co-flowering Erica species from the Cape Floristic Region (South Africa). South African Journal of Botany, 132, 371–377. 10.1016/j.sajb.2020.06.007

96. Vizentin-Bugoni, J., Maruyama, P. K., de Souza, C. S., Ollerton, J., Rech, A. R., & Sazima, M. (2018). Plant-pollinator networks in the tropics: A review. Ecological Networks in the Tropics: An Integrative Overview of Species Interactions from Some of the Most Species-Rich Habitats on Earth, 73–91.

97. Waser, N. M. (1986). Flower Constancy: Definition, Cause, and Measurement. The American Naturalist, 127(5), 593–603. 10.1086/284507

98. Waser, N. M., & Price, M. V. (1985). The effect of nectar guides on pollinator preference: Experimental studies with a montane herb. Oecologia, 67, 121–126.

99. Weiss, M. R. (2001). Vision and learning in some neglected pollinators: Beetles, flies, moths, and butterflies. Cognitive Ecology of Pollination, 171–190.

100. Wells, H., Hill, P. S., & Wells, P. H. (1992). Nectarivore foraging ecology: Rewards differing in sugar types. Ecological Entomology, 17(3), 280–288. 10.1111/j.1365-2311.1992.tb01059.x

101. Wells, H., & Rathore, R. R. (1994). Foraging ecology of the Asian hive bee, *Apis cerane indica*, within artificial flower patches. Journal of Apicultural Research, 33(4), 219–230.

102. Wells, H., & Wells, P. H. (1983). Honey bee foraging ecology: Optimal diet, minimal uncertainty or individual constancy? The Journal of Animal Ecology, 829–836.

103. Wilson, P., & Stine, M. (1996). Floral constancy in bumble bees: Handling efficiency or perceptual conditioning? Oecologia, 106, 493–499.

104. Yoshida, M., Itoh, Y., Ômura, H., Arikawa, K., & Kinoshita, M. (2015). Plant scents modify innate colour preference in foraging swallowtail butterflies. Biology Letters, 11(7), 20150390. 10.1098/rsbl.2015.0390

105. Young, A. M., Kohl, P. L., Rutschmann, B., Steffan-Dewenter, I., Brockmann, A., & Dyer, F. C. (2021). Temporal and spatial foraging patterns of three Asian honey bee species in Bangalore, India. Apidologie, 52(2), 503–523.

106. Yourstone, J., Varadarajan, V., & Olsson, O. (2023). Bumblebee flower constancy and pollen diversity over time. *Behavioral Ecology*, arad028.

